# Chronospaces: an R package for the statistical exploration of divergence times promotes the assessment of methodological sensitivity

**DOI:** 10.1101/2024.02.04.578835

**Authors:** Nicolas Mongiardino Koch, Pablo Milla Carmona

## Abstract

1. Much of our understanding of the history of life hinges upon time calibration, the process of assigning absolute times to cladogenetic events. Bayesian approaches to time scaling phylogenetic trees have dramatically grown in complexity, and depend today upon numerous methodological choices. Arriving at objective justifications for all of these is difficult and time consuming. Thus, divergence times are routinely inferred under only one or a handful of parametric conditions, often times chosen arbitrarily. Progress towards building robust biological timescales necessitate the development of better methods to visualize and quantify the sensitivity of results to these decisions.
2. Here, we present an R package that assists in this endeavor through the use of chronospaces, i.e., graphical representations summarizing variation in the node ages contained in time-calibrated trees. We further test this approach by estimating divergence times for three empirical datasets—spanning widely differing evolutionary timeframes—using the software PhyloBayes.
3. Our results reveal large differences in the impact of many common methodological decisions, with the choice of clock (uncorrelated vs. autocorrelated) and loci having strong effects on inferred ages. Other decisions have comparatively minor consequences, including the use of the computationally intensive site-heterogeneous model CAT-GTR, whose effect might only be discernible for exceedingly old divergences (e.g., the deepest eukaryote nodes).
4. The package *chronospace* implements a range of graphical and analytical tools that assist in the exploration of sensitivity and the prioritization of computational resources in the inference of divergence times.

## INTRODUCTION

The discovery of the “molecular evolutionary clock” by Zuckerkandl and Pauling (1965), who first described the relative constancy in the rates of change of molecular sequences both through time and across clades, is a hallmark of the advent of molecular biology. By establishing a proportionality between the amount of amino acid or nucleotide differences that can be directly observed and the time that must have elapsed since those sequences last shared a common ancestor, the molecular clock greatly expands the insights that can be drawn from phylogenetic trees. As such, scaling phylogenies to absolute (or geologic) time is often the first step to explore macroevolutionary patterns and processes at deep timescales, and one on which modern comparative biology relies on (Pennell & Harmon 2013).

Bayesian methods of divergence time estimation have become highly sophisticated (Kumar & Hedges 2016), being able to accommodate among-lineage rate variation through the use of relaxed clocks (Thorne *et al*. 1998; Drummond *et al*. 2006), exploit diverse sources of both character and temporal information (Ronquist *et al*. 2012; O’Reilly *et al*. 2015), and use complex tree-building models that account for fossilization and sampling rates (Heath *et al*. 2014) and accommodate gene tree discordances (Rannala *et al*. 2020). While this increased complexity has allowed for an improved modelling of diversification, it has also drastically increased the number of methodological choices that the average scientist is confronted with. Dating software require users to specify which among several competing models of diversification, trait evolution, and among-lineage rate variation—the three distinct elements of the tripartite framework of Bayesian dating (Warnock & Wright 2020)—are enforced. Each of these models includes a sizable number of parameters for which prior probability distributions need to be specified. Furthermore, even though progress has been made regarding computational efficiency (Dos Reis & Yang 2019; Douglas *et al*. 2022; Barba-Montoya *et al*. 2023), most methods remain limited in terms of the size of the datasets that can be run within reasonable times, forcing further decisions regarding taxon sampling and locus choice. The choice of number, location, and shape of fossil-derived temporal constraints adds yet another layer of complexity to these analyses. These methodological decisions can have a sizable effect on the inference of Bayesian time-calibrated trees (Warnock *et al*. 2011; Magallón *et al*. 2013; Eme *et al*. 2014; Mongiardino Koch *et al*. 2022; Sauquet *et al*. 2022)—also known as chronograms—producing potentially incongruent reconstructions of biological diversification.

Two broad approaches have been explored to overcome this methodological uncertainty. First, numerous methods have been developed to justify model selection, allowing the identification of a set of parametric conditions considered optimal. These methods, however, have their own limitations: model fit statistics do not reliably discriminate between partition and mixture models (Crotty & Holland 2022), while alternative tree priors might not be amenable to comparison using standard methods such as Bayes factors (May & Rothfels 2023). General purpose approaches, such as cross-validation, are too computationally consuming to be used exhaustively (Zhou *et al*. 2007; Lartillot 2023), while methods designed to tackle specific aspects of the analysis can arrive at opposite conclusions depending on unassessed decisions (e.g., detecting autocorrelated rates can depend on which model of substitution is used to infer the tree being evaluated; Mongiardino Koch *et al*. 2022). The alternative to the use of model selection algorithms involves sensitivity analyses, which explore the robustness of results to a range of conditions (e.g., Sanderson & Doyle 2001; Britton *et al*. 2007; Warnock *et al*. 2011; Howard *et al*. 2022; Mongiardino Koch *et al*. 2022). However, there is a dearth of tools that provide efficient ways of summarizing multiple chronograms in either quantitative or visual ways, as well as a lack of criteria for gauging the relative impact that different parameters have on the final results.

Here, we present *chronospace*, an R package (R Core Team 2022) for the statistical exploration of time calibration. The package implements a set of visualization techniques that allow for a better assessment of the sensitivity of divergence times to methodological decisions. It also provides new tools to quantify the overall impact of each individual methodological choice, providing an estimate of its effect size. We illustrate its functionalities by inferring divergence times for three empirical datasets spanning widely differing evolutionary timescales. We do so using the program PhyloBayes v.4.1 (Lartillot *et al*. 2013) which implements the complex site-heterogeneous model CAT-GTR (Lartillot & Philippe 2004).

## MATERIALS AND METHODS

### A) Chronospaces: the approach

Chronospaces are low-dimensional graphical representations that summarize variation in the absolute node ages of populations of chronograms (Mongiardino Koch *et al*. 2022). They are similar to treespaces (Hillis *et al*. 2005) in that they provide a visual summary of populations of phylogenetic trees, but focus exclusively on temporal information. To do so, the chronograms analyzed must share the same set of nodes, representing posterior distributions of Bayesian time-calibrated analyses run under a fixed topology. While this limits their application, a two-step approach in which a preferred topology is first inferred and then time scaled has become common in the phylogenomic literature due to the computational burden of simultaneous inference.

Whereas treespaces are generated from distance matrices that capture topological similarity (Smith 2022)—and are thus representations that are disconnected from the original variables— chronospaces are built using ordination methods that reorganize variation, capturing major patterns in a few synthetic axes. Here, we rely on between-group principal component analysis (bgPCA; Dolédec & Chessel 1987; Dolédec & Chessel 1989; Yendle & MacFie 1989), an alternative to traditional PCA that ordinates the covariance matrix of multivariate means for predefined groups of observations. Within the context of a chronospace, posterior topologies constitute the ‘observations’, and the ages of each of their internal nodes are the ‘variables’ (see also Fig. 1). When analyzing populations of topologies inferred under different analytical conditions (e.g., alternative clock models), these constitute distinct levels of a grouping variable. The estimated bgPCA axes will be those capturing the largest variation in group (i.e., level) means, and onto which individual chronograms are projected. The result is an ordination that maximizes the separation between groups through a rigid rotation of space, preserving information regarding which nodes are more variable in age, and which factors this variation is associated with. The approach will result in *g-1* bgPCA axes, where *g* is the number of groups, with the remainder of multidimensional structure occurring on residual axes. Within the biological literature, bgPCA has been mostly used to explore the discrimination of groups of observations using microarray (Culhane *et al*. 2002) and geometric morphometric (Cardini *et al*. 2019) data. While bgPCA can introduce spurious patterns when the number of variables exceeds the number of observations (Cardini *et al*. 2019), this does not represent an issue within the context of chronospaces as the number of nodes in a phylogenetic tree (i.e., the number of variables) will typically be much lower than the number of trees sampled in a Bayesian posterior distribution (i.e., the number of observations).

**Figure 1:**
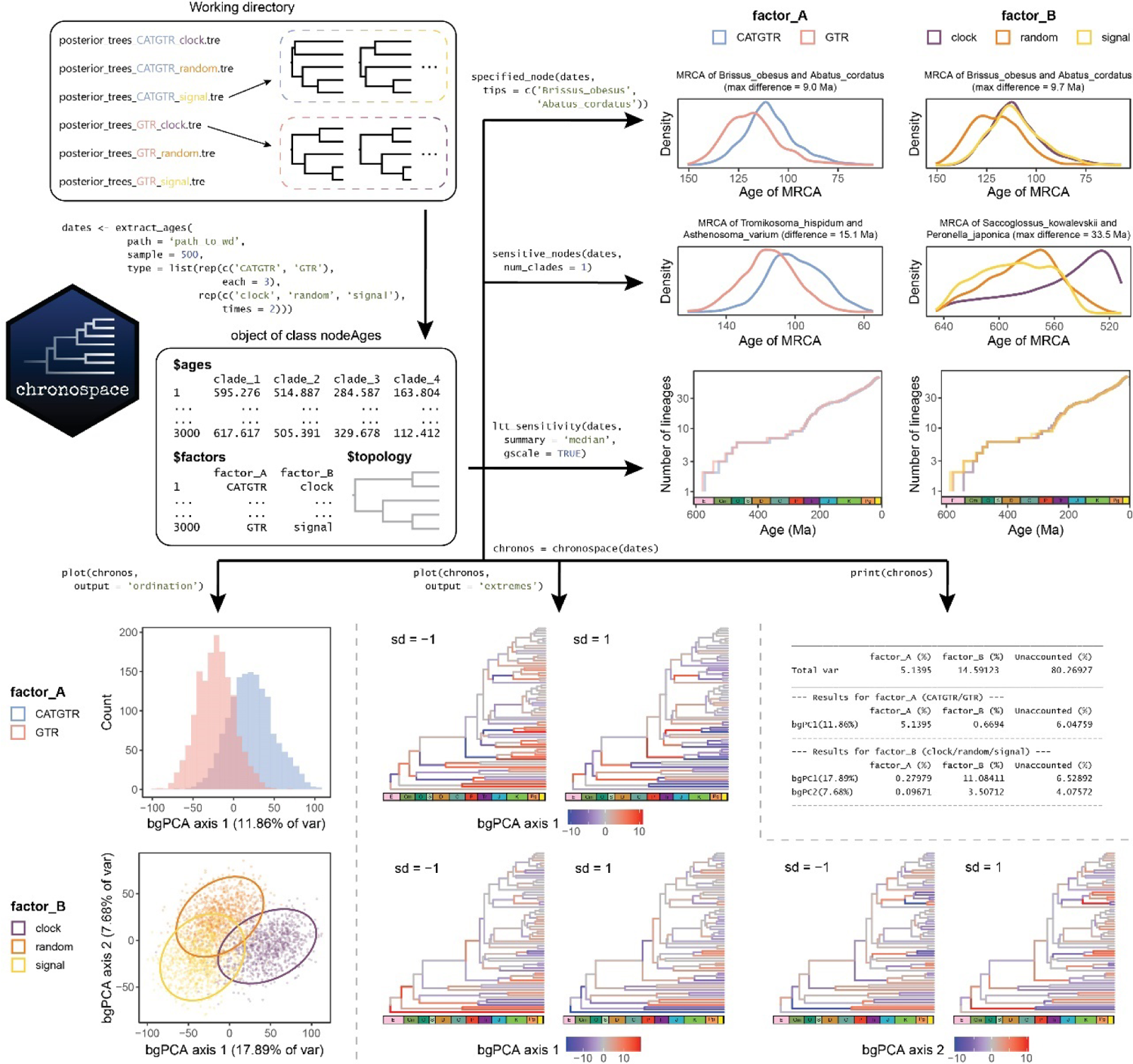
Structure of the R package *chronospace* and analytical workflow, represented using the data that accompanies the package. Populations of posterior topologies from time calibration analyses present in the working directory are imported, and their node ages annotated and stored as an object of class nodeAges using function extract_ages. The package provides functions to visualize the effect induced by analytical conditions on a node of interest (specified_node), and on the overall history of diversification (ltt_sensitivity). It also detects the nodes whose ages change the most between analyses, and plots their posterior distributions (sensitive_nodes). These plots can optionally include a chronostratigraphic scale (shown here in the lineage-through-time plot). Finally, a chronospace object can be built and used to summarize the overall effect of methodological decisions on the results. These can be visualized as bgPCA-summarized chronospaces (which will be univariate or multivariate, depending on the number of levels of each factor) and/or chronogram warping plots. The latter depict the patterns of branch length contraction/expansion that are captured by each bgPCA axis, and are akin to the landmark deformation plots that typically accompany the morphospace axes in a geometric morphometric analysis (hence, their name). All plots are generated with *ggplot2* (Wickham 2016) and several extensions, providing the user with control over basic visualization aspects (object colors, sizes, transparencies, etc.). Chronospace summaries can also be printed in the form of a table reporting the partitioning of variance.

bgPCA axes capture the main directions of change in divergence times that are associated with a given methodological decision. If the chronograms being analyzed are derived from a set of inferences varying multiple parametric conditions, separate bgPCA rotations can be applied to the same chronospace, deriving a set of eigenvectors and eigenvalues for each one of the factors being varied. The proportion of the total variance accounted for by each factor, as well as the proportion that remains unaccounted, is estimated both for the entire chronospace as well as for each individual bgPCA axis by calculating the corresponding sums of squares for the balanced model. Chronospaces thus provide a means of summarizing the entire temporal information of individual chronograms, and exploring the effects (i.e., direction and magnitude of change) associated with alternative dating approaches. This information can be used to aid in the interpretation of the overall sensitivity of divergence times, as well as rank methodological choices from the most to the least impactful.

### B) chronospace: the package

*chronospace* is an R package (R Core Team 2022) available for installation from the GitHub repository (removed for anonymity). It relies on functions from phylogenetic packages *ape* (Paradis & Schliep 2018), *phangorn* (Schliep 2011), and *phytools* (Revell 2012), visualization packages *deeptime* (Gearty 2022), *ggpubr* (Kassambara 2020), *ggtree* (Yu *et al*. 2017), and *patchwork* (Pedersen 2020), as well as data handling functionalities from the *tidyverse* (Wickham 2017).

The range of functions and capabilities implemented in the package are depicted in Figure 1 and further documented on the GitHub repository. The typical workflow would begin with the use of function extract_ages which gathers node ages from posterior distributions of Bayesian timetrees inferred under a constrained topology. Typically, this would include trees from multiple runs (as most of the methods implemented are designed to facilitate the comparison of results obtained under different analytical conditions), but some analyses can be done with a single set of trees. Chronograms are read from files in Newick format (one per run and/or chain). While not all phylogenetic software generate this type of output, trees can be first converted and saved to file using functions from other R packages, such as *treeio* (Wang *et al*. 2020). Function extract_ages can then import the trees, extract and format the node ages contained in them, and annotate the data with a set of factors representing the parameters changed between different runs. The constrained tree topology is also saved in order to generate graphical output. This information (ages, factors, and topology) is stored as an object of class nodeAges. Multiple graphical outputs can be directly obtained from this object: posterior distributions of node ages can be plotted using both sensitive_nodes, which helps identify and visualize the nodes whose inferred ages change the most between analyses, or specified_node, which plots ages for a target node defined by the user; lineage-through-time plots can also be generated using function ltt_sensitivity to depict the overall effect of methodological choices on the inferred history of diversification (including different options for portraying uncertainty). Finally, bgPCA-rotated chronospaces are generated using function chronospace, which creates an object of the same class, and its associated printing and plotting functions summarize the results of the analysis in both graphical and quantitative ways. The remainder of this article focuses on highlighting the use of the latter.

### C) Sensitivity of divergence times across three empirical datasets

Three genome-scale datasets of protein-coding loci were obtained from the literature, designed to tackle the phylogenetic relationships and divergence times of Curculionoidea (65 terminals, 517 loci; Shin *et al*. 2018), Decapoda (90 terminals, 410 loci; Wolfe *et al*. 2019), and Eukaryota (136 terminals, 313 loci; Strassert *et al*. 2021). Studies were selected so as to span widely differing evolutionary time frames (Figures 2). For each of these, we gathered a phylogenomic dataset (including a concatenated matrix coded as amino acids and an associated partition file), the rooted tree topology preferred by the authors, and a list of node age calibrations derived from the fossil record. If studies included multiple outgroups, only those forming part of the sister-group clade to the ingroup were retained. The eukaryote dataset lacked outgroups altogether.

**Figure 2:**
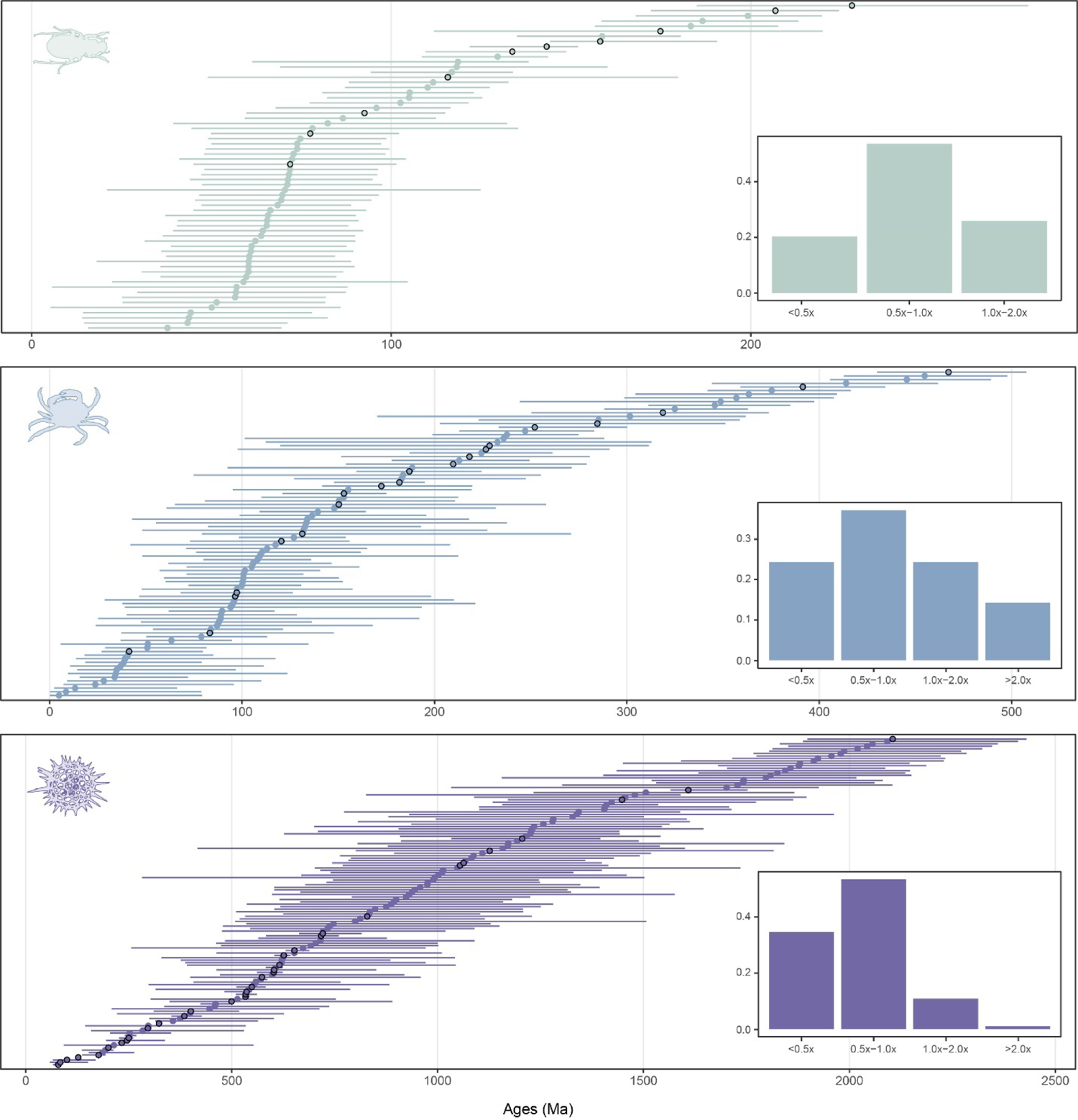
Uncertainty and sensitivity of node age estimates. Values shown are the median and width of pooled 95% CI per node across all analytical conditions. Fossil calibrations were placed on nodes highlighted with black margins. Insets show the distribution of the relative widths of pooled 95% CIs, obtained by dividing absolute widths by the median values. Only unconstrained nodes were tallied.

Divergence times were inferred using PhyloBayes v.4.1 (Lartillot *et al*. 2013) using a constrained tree topology. Four different methodological decisions were assessed: the type of clock, the model of molecular evolution, the prior distribution applied to calibrated nodes, and the approach used to subsample loci. The first three of these are parameters specified when running the program. Divergence times were obtained applying both an autocorrelated log-normal (-ln; Thorne *et al*. 1998) and an uncorrelated gamma (-ugam; Drummond *et al*. 2006) clock; implementing unpartitioned site-homogenous GTR + G (-gtr; Tavaré 1986) or site-heterogenous CAT-GTR + G (-cat -gtr; Lartillot & Philippe 2004) models of molecular evolution; and using prior distributions of node ages that were either uniformly distributed between stratigraphic bounds (-ilb), or implemented a truncated Cauchy minimum bound (-lb; Inoue *et al*. 2010). Default parameter values were used for the latter (*p* = 0.1, *c* = 1), which results in the prior probability peaking at 110% the age of the minimum bound. The tree prior was set to a birth-death process with free parameters and soft calibration bounds were activated (Yang & Rannala 2006), allowing for 5% of prior probability to lie outside of the specified bounds for calibrated nodes (-bd -sb). Default settings were used for all other parameters.

Regarding the fourth and last decision evaluated (the set of loci used), phylogenomic datasets were subsampled using strategies intended to achieve four distinct goals: A) minimize topological conflict; B) improve levels of phylogenetic usefulness; C) maximize the amount of information; and D) enhance clock-like behavior. For the first of these, loci chosen corresponded to those displaying the minimum Robinson-Foulds (RF) distance (Robinson & Foulds 1981) between gene trees and the constrained species tree. For the second, loci were chosen using the multivariate subsampling strategy *genesortR* (Mongiardino Koch 2021; Mongiardino Koch & Thompson 2021), which identifies loci that simultaneously display high phylogenetic signal and low levels of potential sources of bias. Subsampled matrices with maximal amount of information were built from loci with the highest levels of occupancy. Finally, clock-likeness was estimated using the variance of root-to-tip distances displayed by gene trees. In the presence of outgroups, gene trees were rooted ahead of estimation using the node corresponding to the last common ancestor of all ingroup taxa; otherwise, gene trees were midpoint rooted along a random internal edge. This is expected to cause few problems for the curculionoid and decapod datasets for which outgroup occupancy was 98.6% and 81.0%, respectively, but could pose an issue for the eukaryote dataset. To explore this caveat, we estimated the variance of root-to-tip distances for all outgroup-containing gene trees using both outgroup and random midpoint rooting. Correlations between the two were high (Peason’s *ρ* = 0.83 for Curculionoidea and 0.92 for Decapoda, both *p* < 0.001). When used to select the top fifth most clock-like loci, both approaches agreed on 75.5% (Curculionoidea) and 78.8% (Decapoda) of them, further reinforcing that loci displaying clock-like evolution can be recognized in the absence of outgroups. Given the positive relationship between the variance of root-to-tip distances and the overall rate of evolution of genes (Mongiardino Koch *et al*. 2022; see Fig. S1), loci were first sorted by rate (estimated using the overall tree length; Telford *et al*. 2014), and both the fastest- and slowest-evolving 15% of loci were excluded. Loci chosen corresponded to those with minimum root-to-tip variances from among those showing relatively average evolutionary rates. Gene trees necessary to measure these properties were inferred using ParGenes v.1.0.1 (Morel *et al*. 2018), using models that minimized the Bayesian Information Criterion and estimating support with 100 bootstrap replicates. A fifth subsampled dataset was derived by selecting loci at random, providing a means of estimating the overall effect of targeted subsampling schemes. To standardize these different matrices in terms of the overall amount of data contained, the loci necessary to first exceed a total length of 10,000 positions were concatenated.

Dating was performed under all combinations of these four methodological decisions, resulting in 40 conditions per dataset, each of which was run using two independent chains for at least 20,000 generations. Posterior topologies were obtained using the verbose output option of the program readdiv, setting a 50% burn-in and thinning chains to every two generations (-x 10000 2 -v), for a total of 5,000 posterior trees per chain. Convergence was visually assessed using log-likelihood trace plots (Fig. S2) and comparing median posterior estimates of node ages and model parameters between replicate chains (Figs. S3-S6). After confirming convergence was attained, time-calibrated trees from separate chains were combined into a single file. The remainder of analyses were performed using package *chronospace*, summarizing each run using a random sample of 500 posterior trees. R code to reproduce each of these steps is available as Supplementary Material S1.

## RESULTS

Figure 2 summarizes the results obtained across the three empirical datasets used, showing the median and 95% confidence interval (CI) of each inferred node age after combining the posterior distributions of individual runs. Producing such overall summaries of temporal information across populations of chronograms is straightforward once the data has been gathered and formatted using the function extract_ages. For the ancient divergences estimated in the eukaryote dataset, pooled 95% CI can span upwards of 1,000 Ma, as is the case for example of the last common ancestor of Amebozoa (median: 1,291.2 Ma, 95% CI: 701.2-1,841.2 Ma). The temporal uncertainty associated with younger divergences is much narrower in absolute terms, yet it can be even larger in relative terms (i.e., after dividing the 95% CI width by its median value). For example, a handful of divergences between closely related achelate decapods show 95% CI widths that are six to seven times the value of the median estimate, with plausible dates spanning the last 142 Ma and falling anywhere between the Quaternary and Early Cretaceous.

The spread of posterior dates shown in Figure 2 is a function of the width of the individual CI of each run (uncertainty) and the degree of overlap of the CIs obtained under different conditions (sensitivity). Chronospaces can help disentangle the two by visualizing and quantifying the contribution of each individual methodological choice to overall levels of sensitivity (Figure 3). In the case of the empirical datasets analyzed, this approach reveals that: A) there is a remarkable difference (over 1,000x) in the overall impact of the parameters explored; and, B) the type of molecular clock and the subsample of loci have larger effect sizes for the all three datasets. The implementation of the complex site-heterogeneous CAT-GTR model seems to have a strong effect only for the eukaryote dataset. The most sensitive nodes per dataset to each of the four factors evaluated are found in Figures S7-S9.

**Figure 3:**
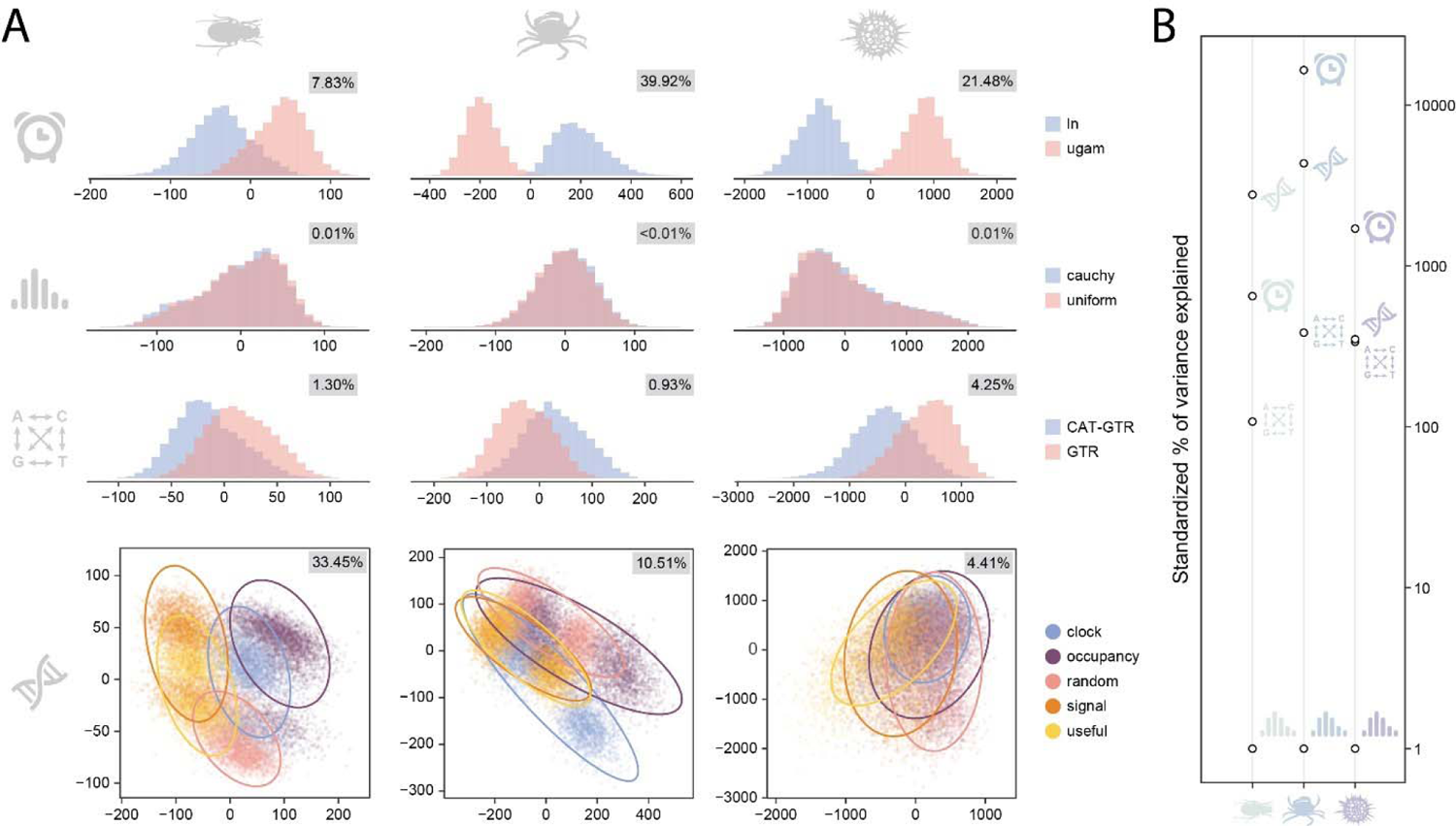
Sensitivity of divergence time estimates to methodological choices. *A.* bgPCA-summarized chronospaces for each factor, from top to bottom: type of molecular clock, prior distribution on constrained nodes, model of molecular evolution, and gene subsample. The proportion of total variance explained by each factor is shown in grey boxes. For the loci subsample, only the first two axes (of four) are shown. Ellipses include 95% of chronograms. *B.* Proportion of total variance explained by each factor, divided by that of the least important factor for each dataset. Axis is log-scaled.

Regarding loci subsampling, chronograms built using high signal and high usefulness loci were very similar across datasets, defining one extreme of the first (major) bgPCA axis (Fig. 4A). The other end of this axis was always populated by chronograms derived from high occupancy loci, and occasionally (for example, in the case of the eukaryote dataset) also by random and clock-like schemes. The branch-length warping plots shown in Figure 4B reveal that the selection of high signal/usefulness loci results in systematically longer internal branches and shorter terminal ones.

**Figure 4:**
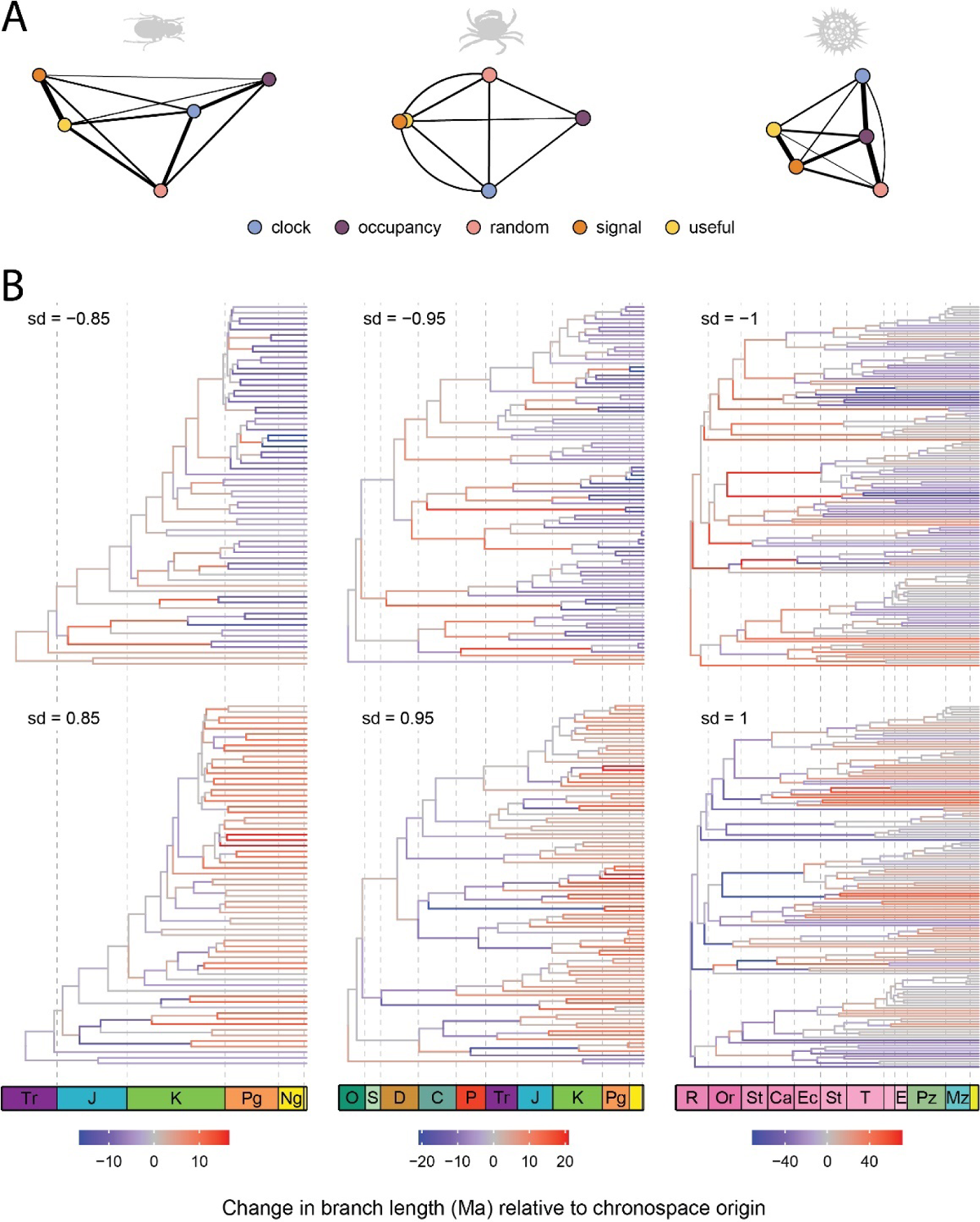
Effect of locus choice on inferred ages. *A.* Chronospaces built discriminating by loci subsampling strategy, as also shown in Fig. 3A. Circles depict the centroids of each distribution; the thickness of the lines connecting them is scaled to the inverse of the Euclidean distances separating them. Locations of centroids are shown on the first two bgPCA axes only, distances were calculated using all dimensions. *B.* Graphical representation of chronograms representing negative (top) and positive (bottom) extremes of bgPC1. The expansion/contraction of branches relative to a chronogram at the origin of bgPC1 is color coded (red/blue, respectively). sd = standard deviation along bgPCA 1; values < 1 are automatically selected by the package in order to avoid generating negative branch lengths. Chronogram warpings induced by other factors are shown in Figures S10-S12.

## DISCUSSION

Understanding evolutionary processes across geologic timescales cannot be done without first establishing an accurate timeline of biological diversification. While phylogenomic datasets, and methods able to handle them, have allowed us to tackle phylogenetic questions that were once intractable (e.g., Philippe *et al*. 1994; Poe & Chubb 2004), they have also introduced a plethora of methodological steps and decisions with downstream effects that we are only beginning to explore (Tan *et al*. 2015; Li *et al*. 2021; Mongiardino Koch 2021; Lozano-Fernandez 2022; Foster *et al*. 2023). In the context of modern Bayesian relaxed-clock dating, these decisions relate to the selection of one among several models that determine how the tree is built, how the characters evolve, and how a clock is enforced—adding up to dozens of parameters that require user-specified prior probability distributions (Álvarez-Carretero & dos Reis 2020; Warnock & Wright 2020). Further decisions need to be taken regarding the character information that is employed, the taxa that are sampled, and the temporal constraints that are enforced. These choices can have strong consequences: they can determine whether we think clades survived mass extinctions (Budd & Mann 2023), emerged through sudden bursts of innovation (Dos Reis *et al*. 2015), participated in events of co-diversification with mutualistic partners (Barba-Montoya *et al*. 2018), and diverged—or not—as a direct consequence of continental drift (Tarver *et al*. 2016).

Most of these methodological decisions are also hard to justify. Coupled with the high computational cost of analyses, empirical evolutionary timescales are generally inferred using one (or a few) among hundreds of possible parametric conditions, often times chosen arbitrarily. A literal reading of the literature assessing the sensitivity of results to methodological factors suggests that every one of these decisions can have large impacts on inferred dates (Hugall *et al*. 2007; Inoue *et al*. 2010; Clark et al. 2011; Duchêne *et al*. 2014; Hirt *et al*. 2017; Barba-Montoya *et al*. 2018; Dos Reis *et al*. 2018; Smith *et al*. 2018; Carruthers & Scotland 2021; Chen *et al*. 2021), providing little *a priori* guidance regarding which should be prioritized given limited computational resources. These assessments are likely also compounded by the choice of dating software, as estimates of sensitivity might be driven by the range of options available, their implementation, or even their default parameterization (which is not always tuned, as is the case of the present study). Progress in pinpointing the magnitude of these effects has been hampered by a general focus on a handful of nodes of interest rather than on tree-wide effects, a scarcity of methods of visualization and summary, as well as of metrics that quantify the effect size of individual decisions in a manner that is comparable, and a lack of testing of whether the relevance of these choices scales with temporal depth.

Chronospaces provide novel ways of visualizing, quantifying, and exploring the sensitivity of divergence time estimates, contributing to the inference of more robust evolutionary timescales (Mongiardino Koch *et al*. 2022; Song *et al*. 2023; Wolfe *et al*. 2023). By representing chronograms as collections of node ages, standard multivariate statistical approaches can be readily employed on populations of Bayesian posterior timetrees. This representation allows for more nuanced visualizations that can effectively convey the overall impact of different choices (Fig. 3A), the relative similarity of trees obtained across runs (Fig. 4A), and the patterns of branch length warping that are induced (Fig. 4B). At the same time, partitioning the total variance into proportions explained by discrete factors using sums of squares can result in tree-wide estimates of sensitivity with which to rank methodological choices by their relative importance, helping prioritize computational resources. While the current implementation is limited to analyses run under a constrained tree topology, chronospaces could potentially be built using the subset of ages derived from nodes present in all (or most) of the posterior trees of a simultaneous-inference analysis.

As shown in Fig. 3B, some decisions affect our reconstruction of diversification histories much more than others, with effect sizes that vary by three to four orders of magnitude. This extreme heterogeneity can help redirect the allocation of computational effort. For example, our results suggest that empirical studies should either carefully validate their choice of evolutionary clock, or run inferences that vary this component of the analysis, before deriving evolutionary insights (as shown previously in both simulated and empirical studies; see Ho *et al*. 2005; Duchêne *et al*. 2014; Delsuc *et al*. 2018; Dos Reis *et al*. 2018; Mongiardino Koch *et al*. 2022; Song *et al*. 2023). Other factors can be arbitrarily fixed without significant consequences, as is the case of the shape of prior distributions on calibrated nodes (as noted previously by Betts *et al*. 2018, although see Clarke *et al*. 2011). We note that the lack of an effect detected here applies exclusively to the use of the Cauchy lower bound (Inoue *et al*. 2010) as implemented in PhyloBayes under default settings, and does not mean that the fossil calibration scheme is of little relevance—in fact, ample evidence points to the opposite (e.g., Near *et al*. 2005; Magallón *et al*. 2013; Montagna *et al*. 2019). The use of prior distributions with other alternative shapes, the enforcement of hard or soft bounds, the choice of different calibration points (as well as their overall number and relative depths), are all major components of dating analyses that were not evaluated here. Similarly, the software of choice can impart idiosyncratic effects (Bapst *et al*. 2016), meaning that all conclusions drawn in this study could also be software-specific. The main rationale for choosing PhyloBayes is that it remains the only dating software implementing the site-heterogeneous CAT-GTR model (Lartillot & Philippe 2004). Despite ongoing controversies, this model holds preeminence in the inference of phylogenomic topologies (Whelan & Halanych 2017; Li et al. 2021; Bujaki & Rodrigue 2022; Al Jewari & Baldauf 2023; Szánthó *et al*. 2023), yet its impact on dating had so far remained unexplored. Our results suggest that the CAT-GTR model induces relatively minor effects. Such robustness of divergence times to the choice of substitution model has been reported before (Du *et al*. 2019; Tao *et al*. 2020; Mongiardino Koch *et al*. 2022), implying that the cost of using computationally intensive models is not outweighed by the benefits. Notably, the relatively high effect of the choice of loci and clock are not affected by the timeframe of the phylogenetic question, and affect equally nodes of Cenozoic age as well as divergences spanning the Proterozoic (Fig. 3B). Only two differences were observed as deeper timescales are probed: there is a decrease in the impact of the loci subsample coupled with an increase in that of the clock model (with both remaining the two most important factors overall); and the choice of substitution model becomes slightly more relevant for extremely ancient (e.g., Precambrian) divergences. The generality of these effects remains to be tested with a broader sample of datasets.

Loci subsampling has become a standard approach in phylogenomic dating, yet again, the effects introduced by the each of numerous available approaches have not been fully characterized. While some authors have advocated for the selection of clock-like genes in the hopes that reduced lineage-specific rate variation will improve model fit and diminish bias, others have favored loci with less topological conflict, as these are expected to harbor higher phylogenetic signal and be less affected by incomplete lineage sorting (ILS). Simulations have confirmed that both of these subsampling schemes result in more accurate divergence times (Smith *et al*. 2018; Carruthers *et al*. 2020; Carruthers *et al*. 2022). Other sensible options include selecting genes based on phylogenetic usefulness (as estimated using *genesortR*; Mongiardino Koch 2021), which should balance both aforementioned strategies by discovering loci with increased clock-likeness and decreased topological incongruence; and maximizing occupancy, increasing the amount of information contained within a matrix of a given size.

Our results show that locus choice is a major factor determining the divergence times obtained, and that alternative schemes impart characteristic signatures on inferred chronograms. Topologically congruent loci (targeted by both the signal and usefulness schemes) produce shorter terminal branches and longer internal ones (particularly those subtending species pairs; Fig. 4B). The opposite effect is induced by maximizing occupancy, and to some extent also by favoring clock-like loci. These same patterns of branch length warping were found in the simulation study of Carruthers *et al*. (2022), in which the selection of congruent topologies was tied to an expansion of internal branch lengths and a reduction of terminal ones. This is in fact the expected behavior if topological incongruence is driven by biological factors such as ILS (Mendes & Hahn 2016). While our analyses were not designed to test this phenomenon, the results are consistent with ILS driving topological incongruence even at exceedingly ancient evolutionary timescales. Even if targeting clock-like sequences can improve the modelling of rate variation, doing so in a manner that does not also account for topological incongruence can result in the inference of distorted evolutionary timescales.

## SUPPLEMENTARY MATERIAL

**Figure S1:**
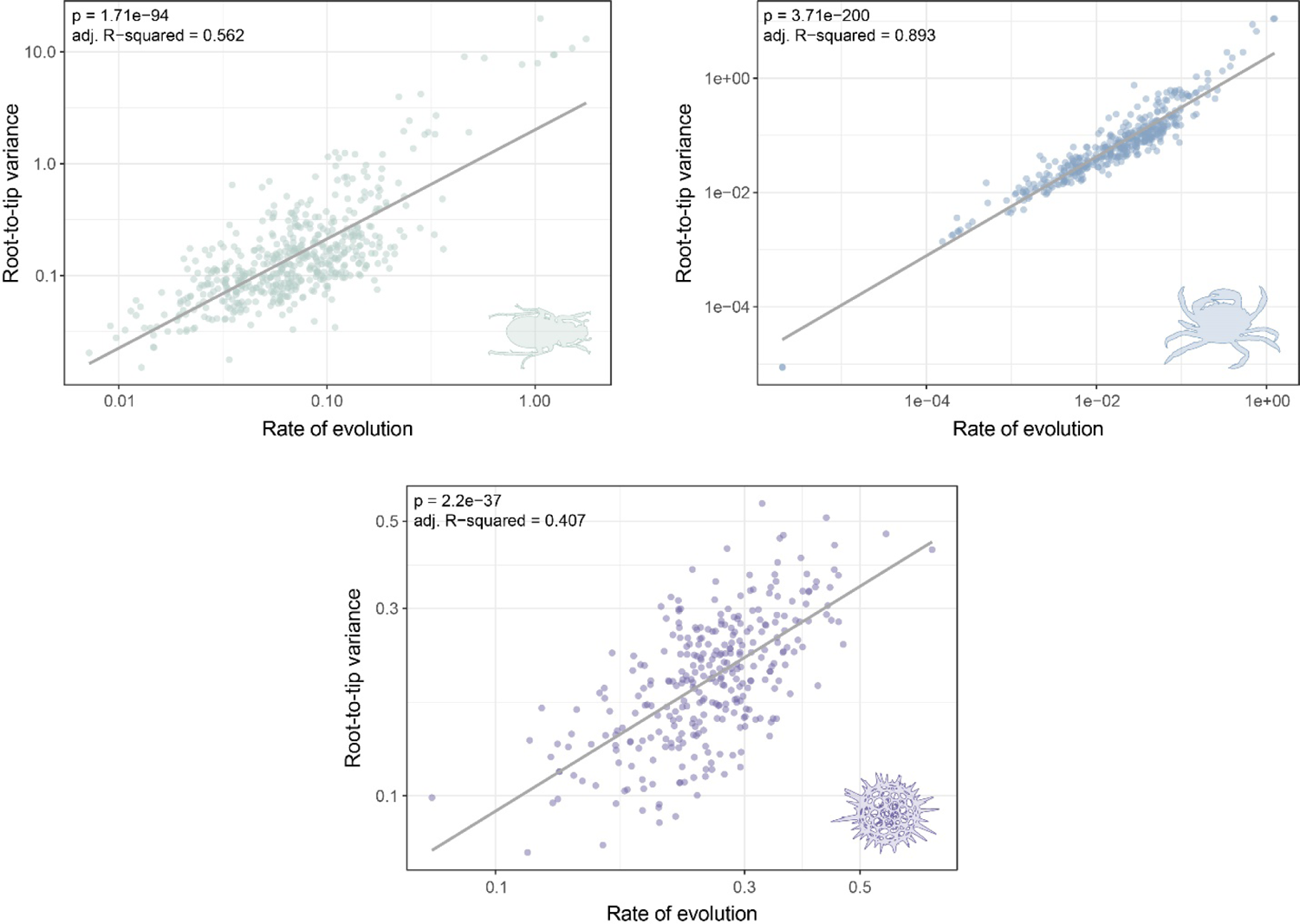
Root-to-tip variance (a proxy for clock-likeness) is strongly determined by the evolutionary rate of loci. Minimizing this variable, a common approach when subsampling matrices for time calibration, also results in selecting genes that evolve at minimum rates. To avoid this confounding effect, clock-like genes from these three matrices were selected after filtering the fastest and slowest evolving 15% of loci. Lines correspond to linear regressions; both axes are log-scaled.

**Figure S2:**
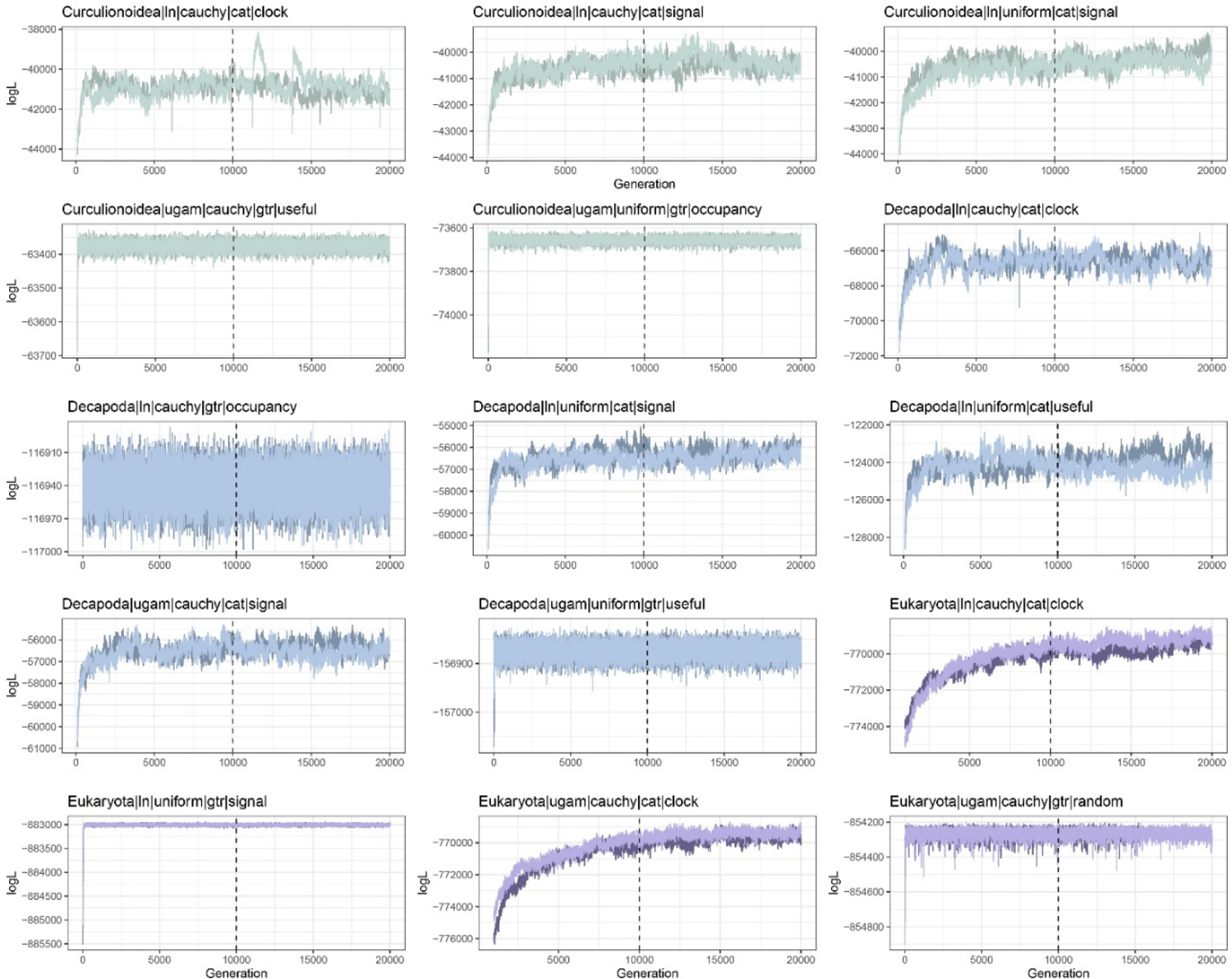
Trace plots of the log-likelihood (logL) of chains. Results correspond to fifteen randomly selected analyses. Dashed lines correspond to the burn-in threshold imposed (10,000 generations). Convergence and stationarity vary between conditions and datasets, with mixing being more challenging when using the site-heterogeneous model CAT + GTR + G, and convergence taking longer for the eukaryote dataset. Nonetheless, both chains of each analysis seem to have converged to the same high posterior region of parameter space by the time the burn-in generations have elapsed. Note that the first few logL values were discarded in order to arrive at a meaningful scaling of the y-axis. The number of excluded generations was 10 for analyses under the GTR + G model, and either 100 (Curculionoidea, Decapoda) or 1,000 (Eukaryota) for analyses under the CAT + GTR + G model. Surprisingly, runs under GTR + G converged within the first 50 generations (sometimes, as is the case of the second decapod plot, within less than 10).

**Figure S3:**
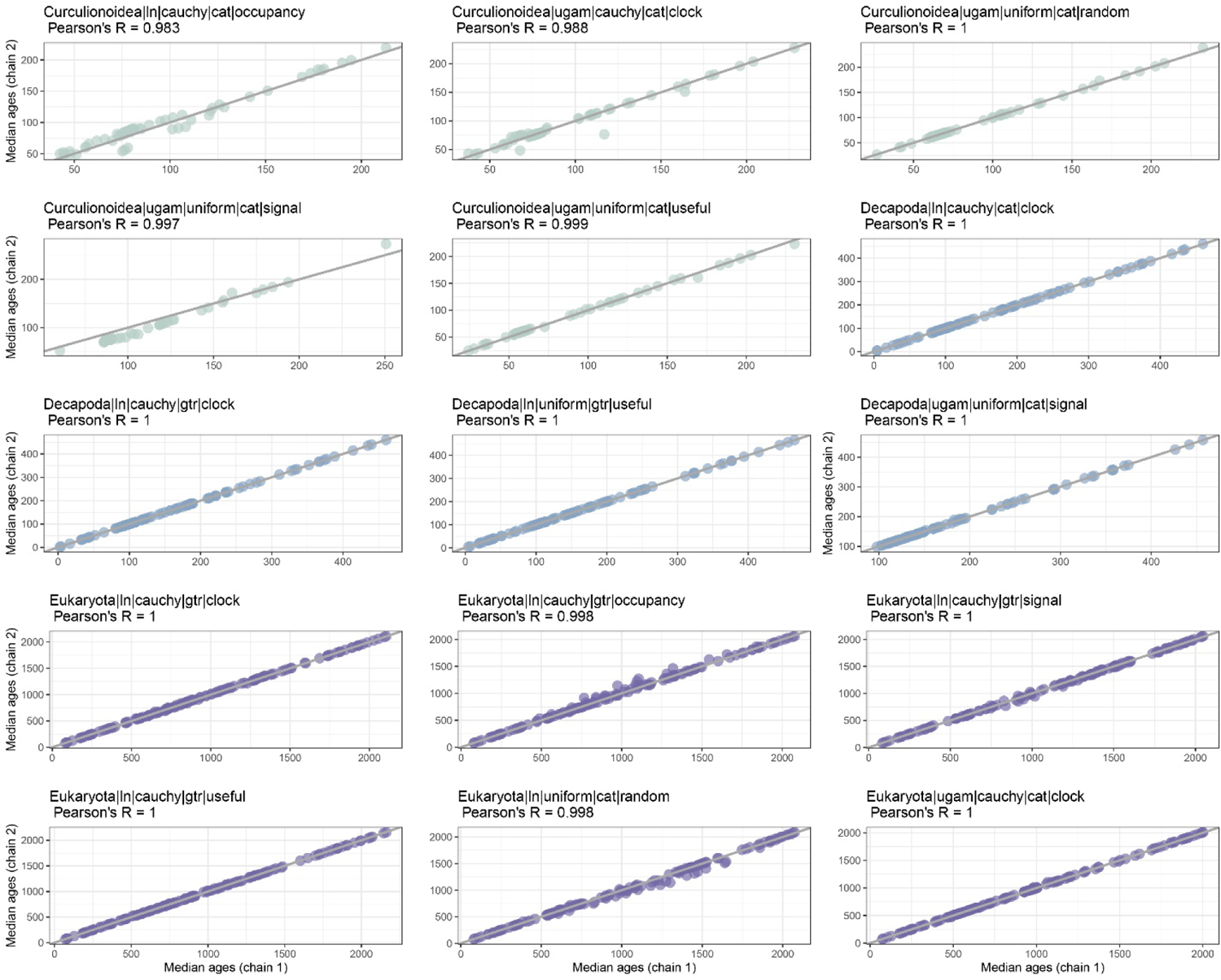
Median node ages for pairs of chains showing appropriate levels of convergence. Results correspond to fifteen randomly selected analyses. Grey lines show 1-to-1 lines. The minimum Pearson correlation coefficient (R) found across all analyses was 0.973.

**Figure S4:**
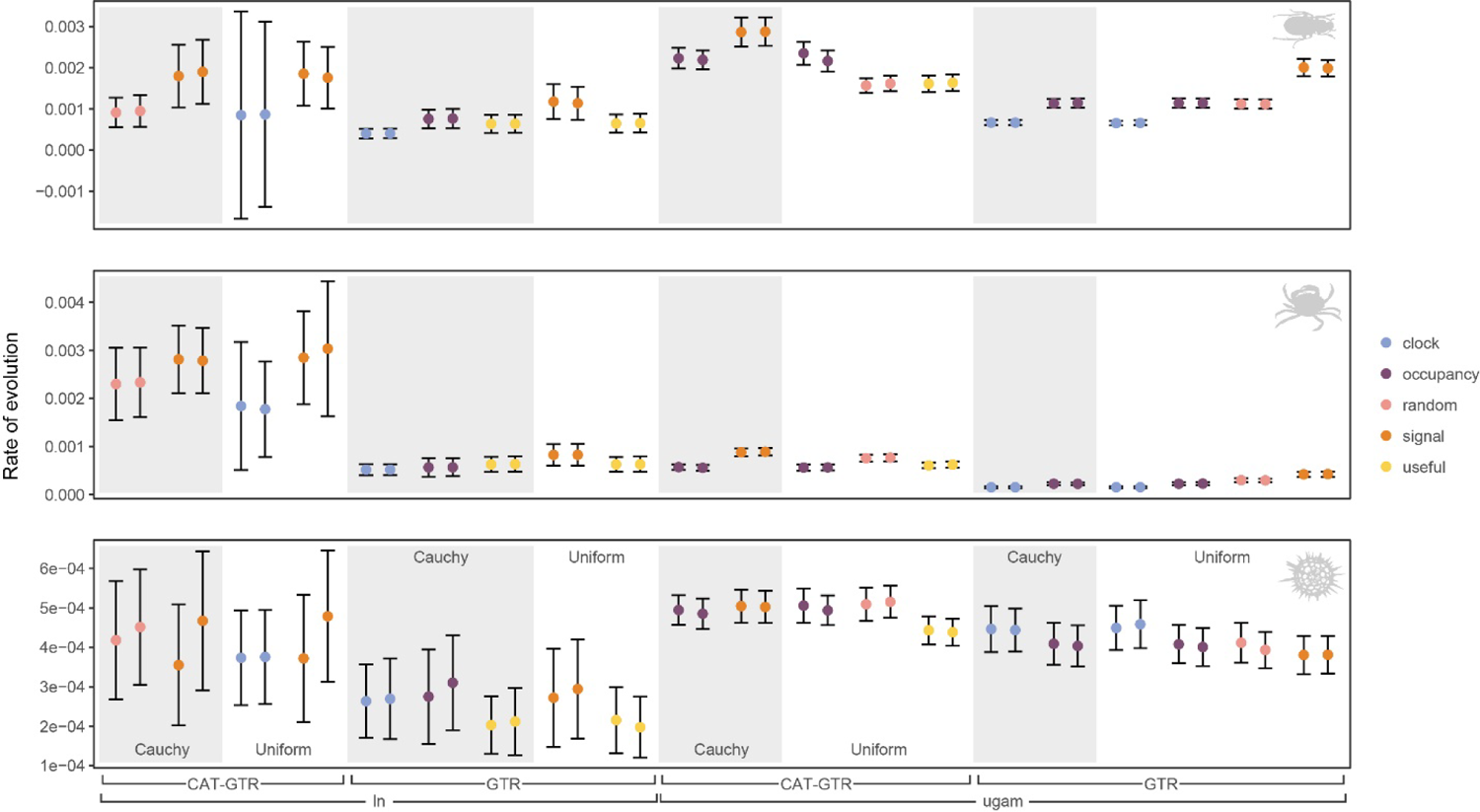
Comparison of the posterior estimates of evolutionary rates across a random sample of 20 analyses per dataset (i.e., 60 analyses total, 2 chains each). Pairs of chains are shown next to each other. Estimates correspond to posterior means ± 1 standard deviation.

**Figure S5:**
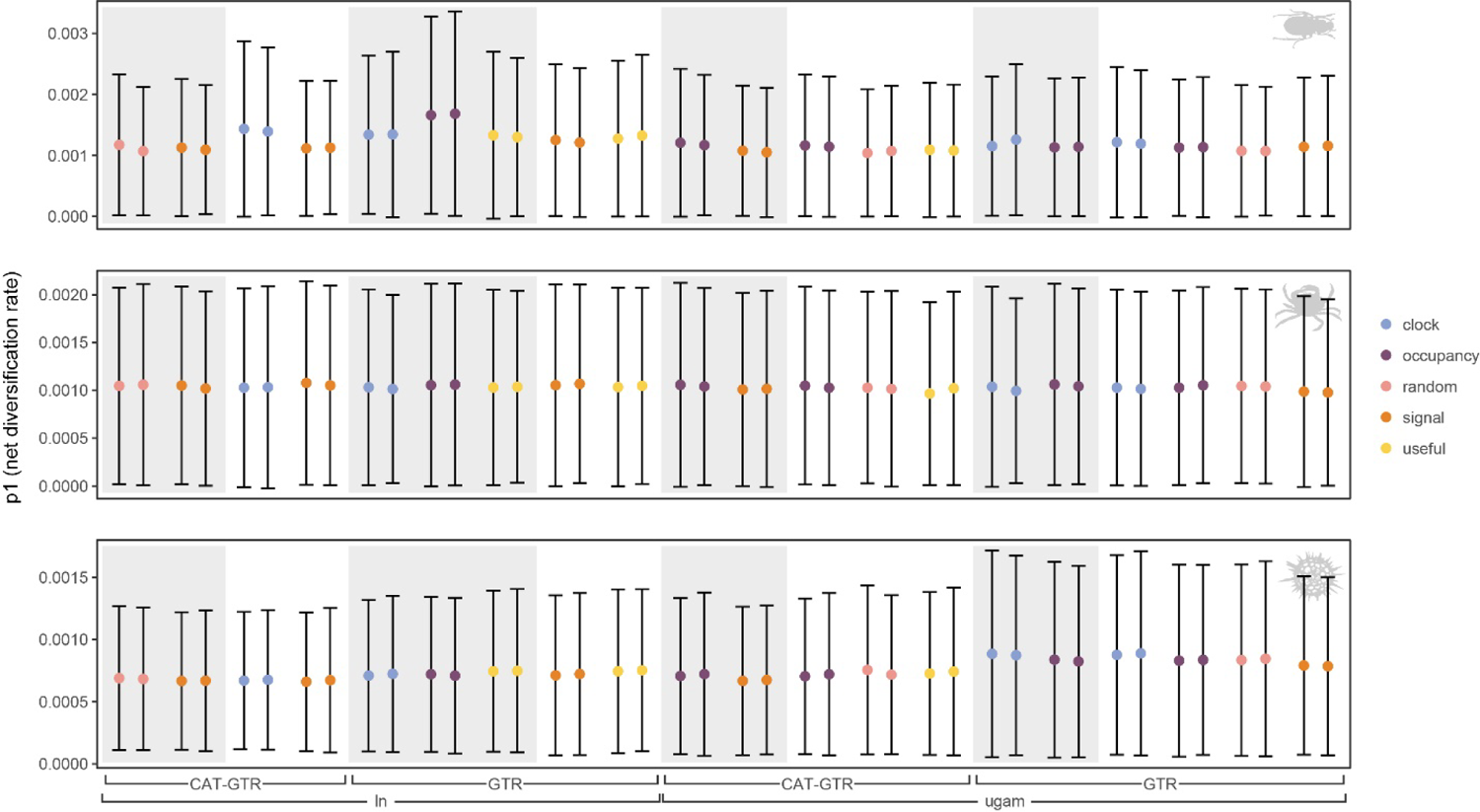
Comparison of the posterior estimates of p1, one of the two parameters of the birth-death tree prior, across a random sample of 20 analyses per dataset (i.e., 60 analyses total, 2 chains each). Pairs of chains are shown next to each other. Estimates correspond to posterior means ± 1 standard deviation. p1 is corresponds to the birth rate (λ) minus the death rate (μ), and is thus the net diversification rate. Data is for the same analyses shown in Fig. S4.

**Figure S6:**
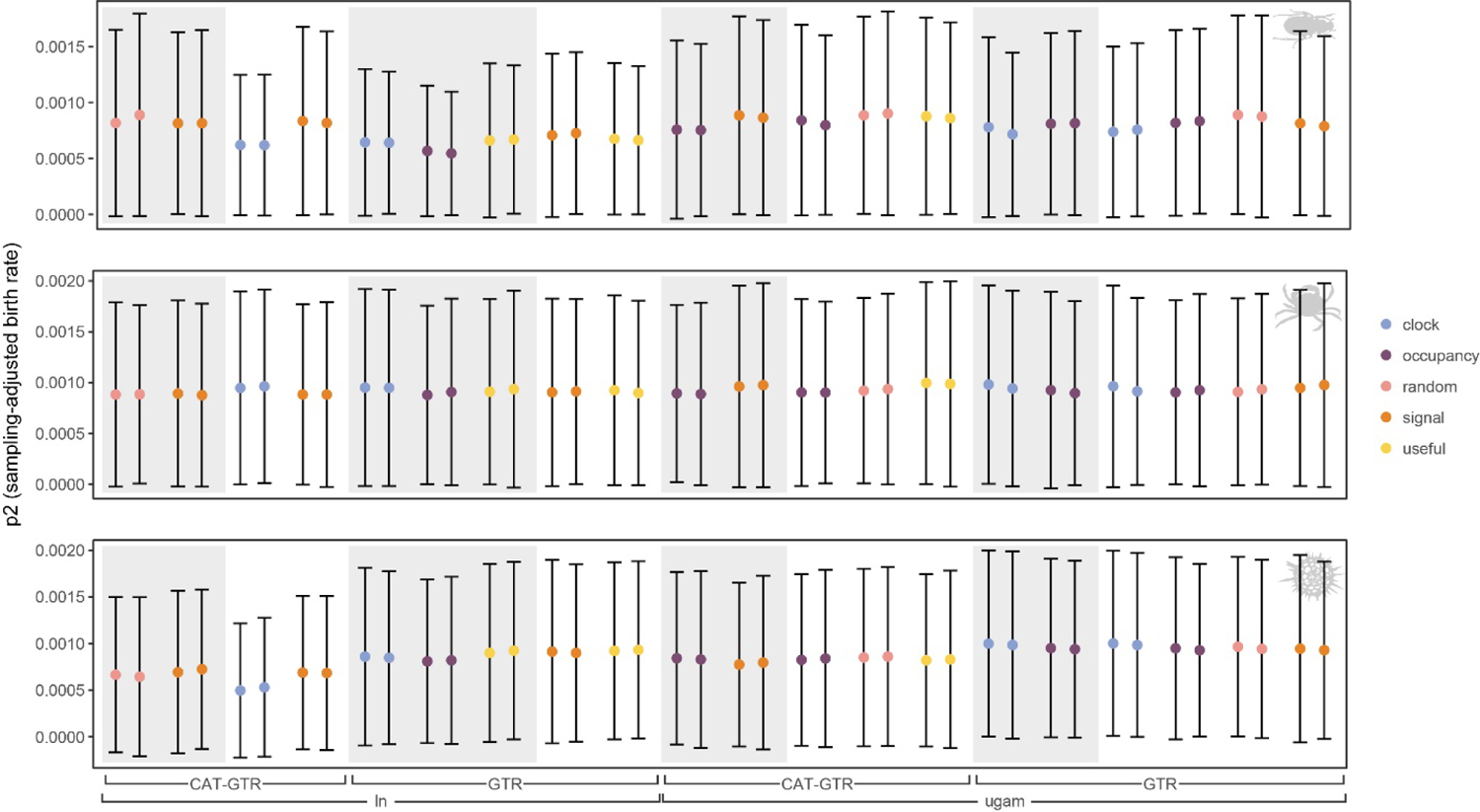
Comparison of the posterior estimates of p2, one of the two parameters of the birth-death tree prior, across a random sample of 20 analyses per dataset (i.e., 60 analyses total, 2 chains each). Pairs of chains are shown next to each other. Estimates correspond to posterior means ± 1 standard deviation. p2 is corresponds to the birth rate (λ) multiplied by the sampling fraction (ρ). Data is for the same analyses shown in Fig. S4.

**Figure S7:**
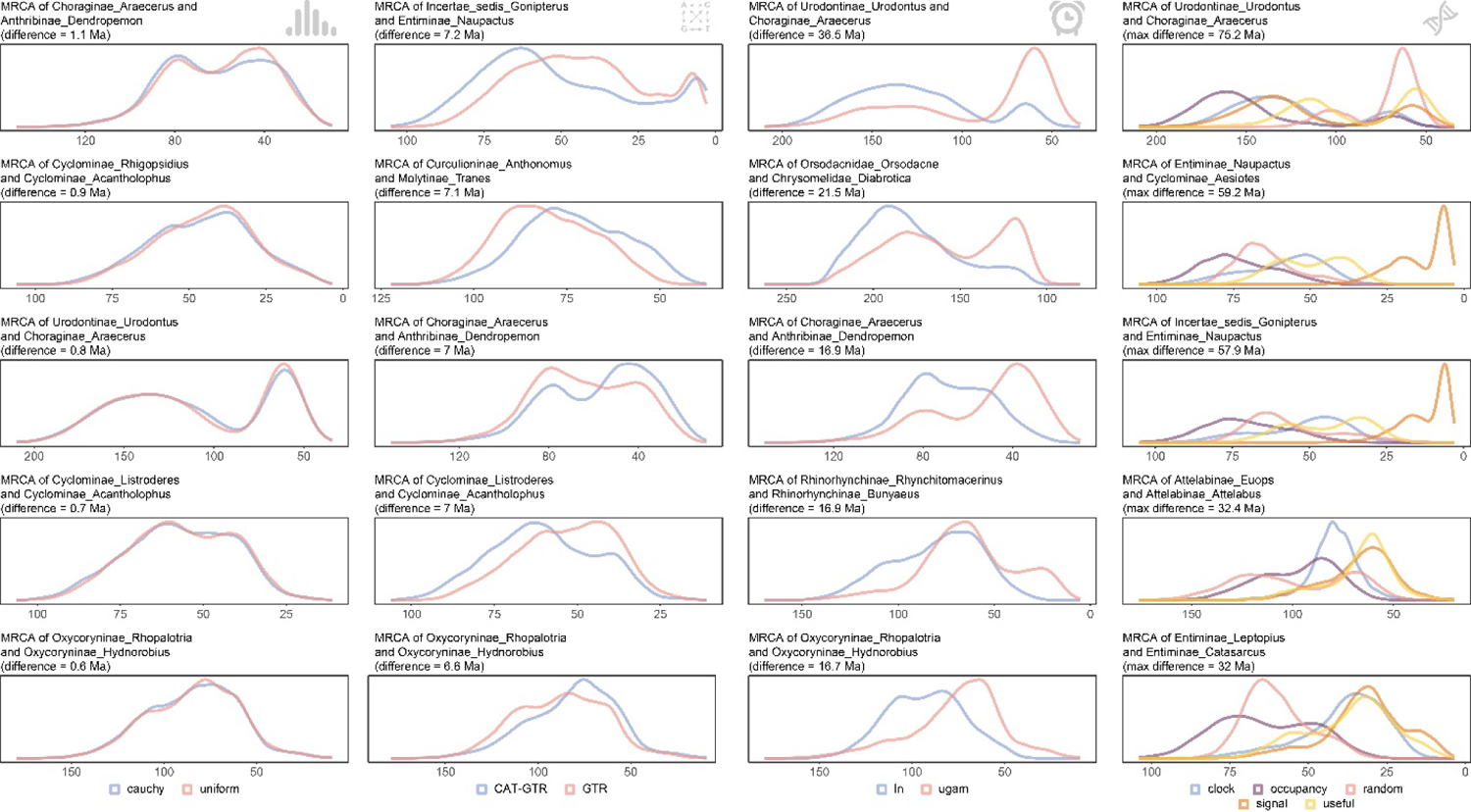
Posterior distributions of ages for the most sensitive nodes of the curculionid dataset. For each methodological choice being varied, the top five most sensitive nodes are depicted. Factors are ordered from the least impactful on the left, to the most impactful on the right (according to the proportions of explained variance, see Fig. 3). Nodes are identified using two randomly-selected terminals from each descendant clade.

**Figure S8:**
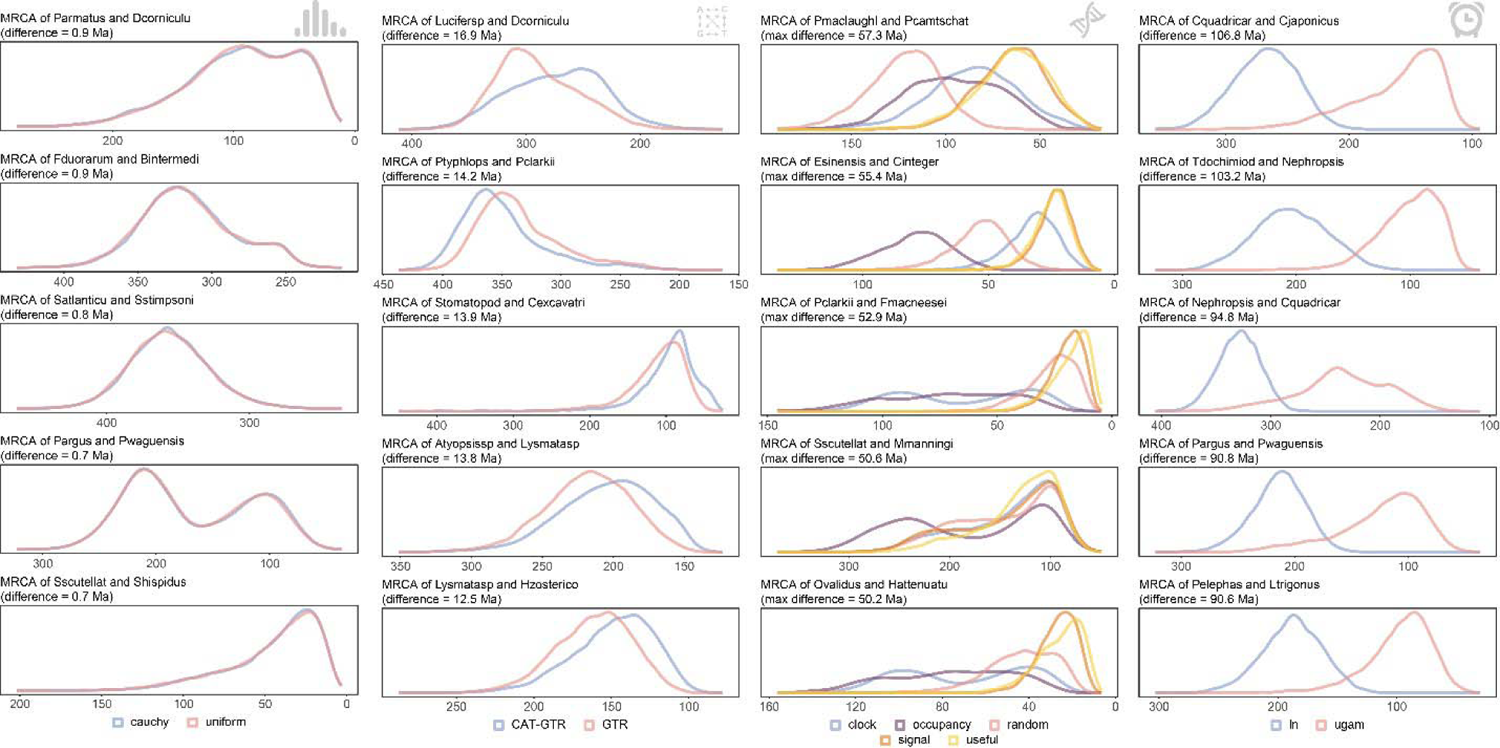
Posterior distributions of ages for the most sensitive nodes of the decapod dataset. For each methodological choice being varied, the top five most sensitive nodes are depicted. Factors are ordered from the least impactful on the left, to the most impactful on the right (according to the proportions of explained variance, see Fig. 3). Nodes are identified using two randomly-selected terminals from each descendant clade.

**Figure S9:**
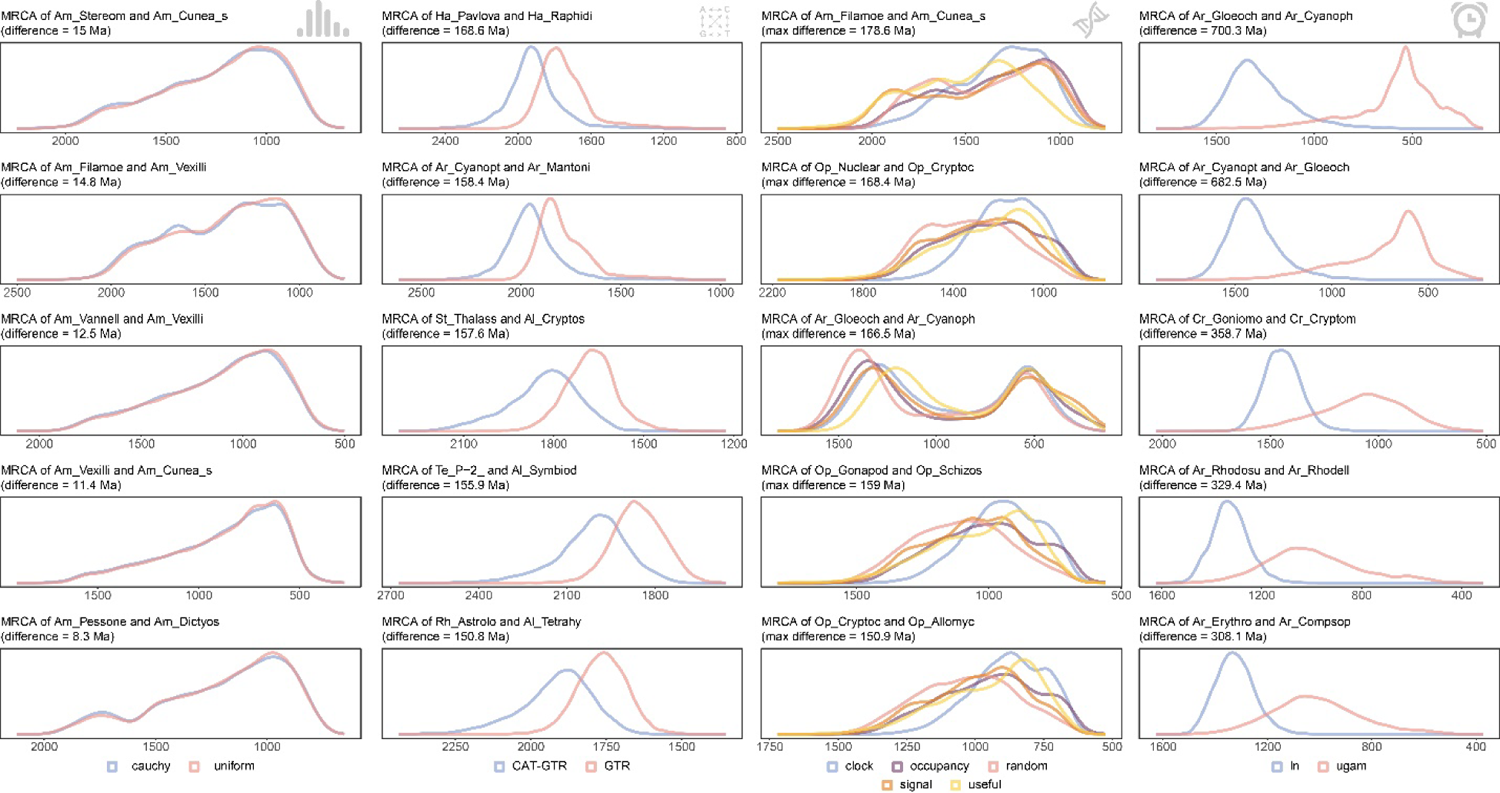
Posterior distributions of ages for the most sensitive nodes of the eukaryote dataset. For each methodological choice being varied, the top five most sensitive nodes are depicted. Factors are ordered from the least impactful on the left, to the most impactful on the right (according to the proportions of explained variance, see Fig. 3). Nodes are identified using two randomly-selected terminals from each descendant clade.

**Figure S10:**
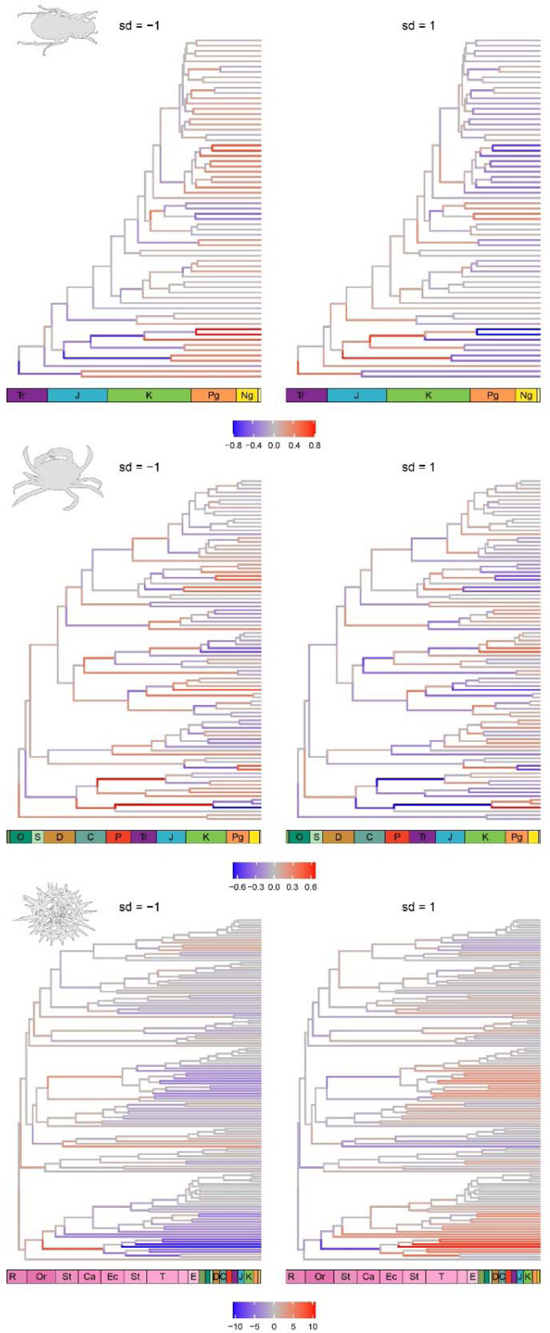
Branch length warping along the bgPCA maximizing the discrimination of chronograms inferred using different node age prior distributions (i.e., the least impactful decision across all datasets; see Fig. 3). This decision seems to induce heterogeneous effects across subclades.

**Figure S11:**
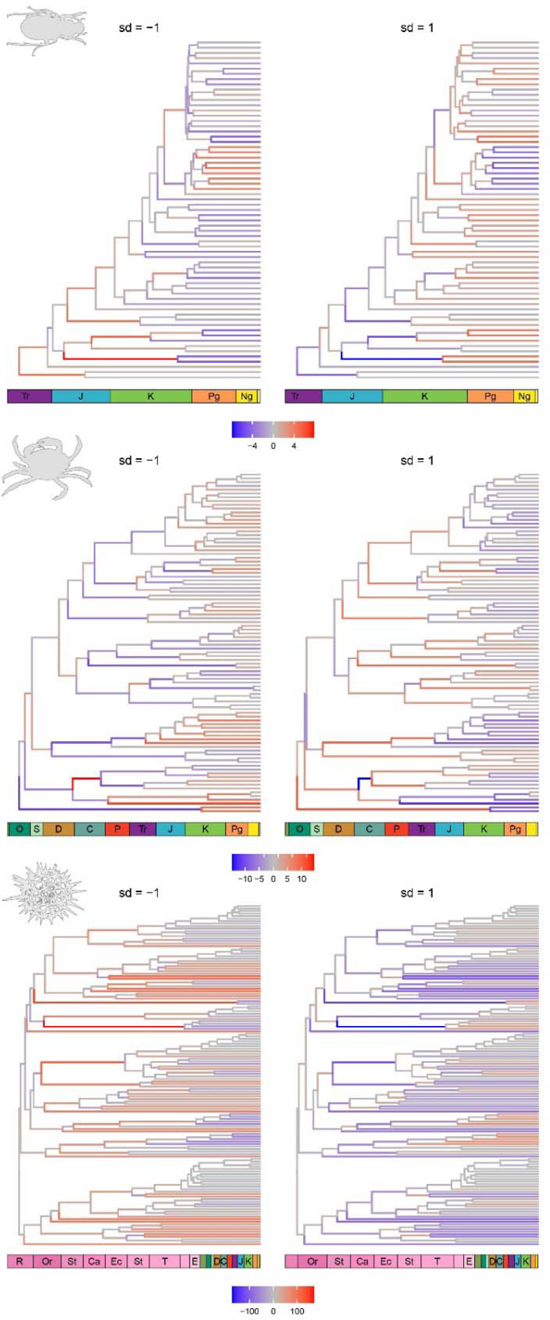
Branch length warping along the bgPCA maximizing the discrimination of chronograms inferred using different models of molecular evolution (i.e., the second least impactful decision across all datasets; see Fig. 3). This decision seems to induce heterogeneous effects across subclades.

**Figure S12:**
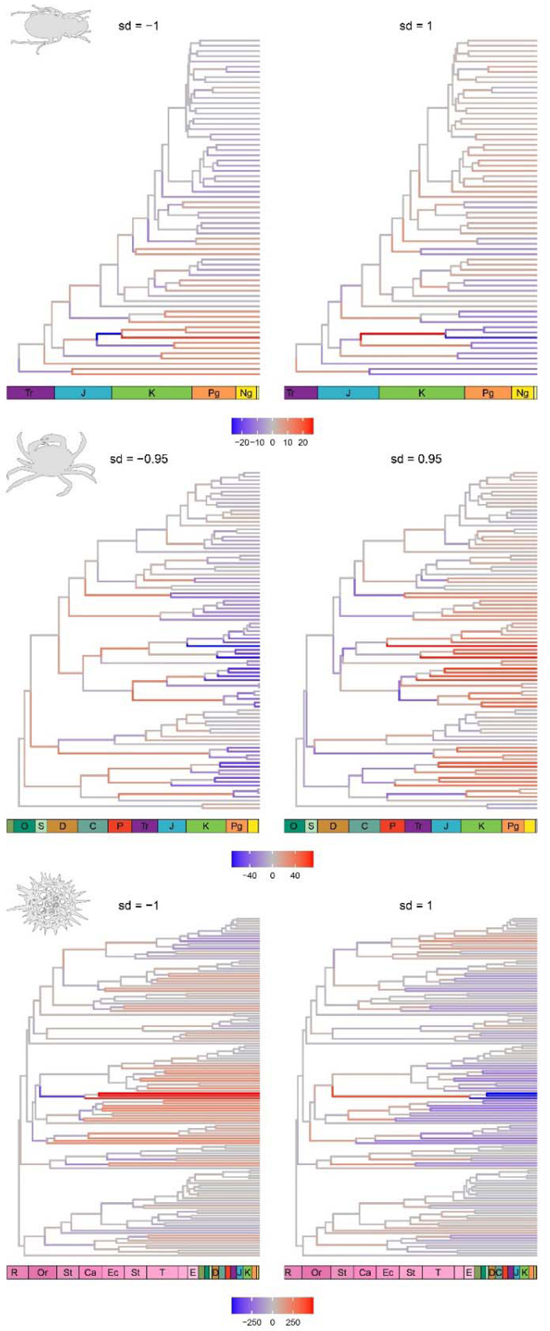
Branch length warping along the bgPCA maximizing the discrimination of chronograms inferred using different clock models (i.e., the first or second most impactful decision across all datasets; see Fig. 3). This decision seems to induce effects that are strong, yet clade specific.

